# On the role of transcription in positioning nucleosomes

**DOI:** 10.1101/2020.04.07.029892

**Authors:** Zhongling Jiang, Bin Zhang

## Abstract

Nucleosome positioning is crucial for the genome’s function. Though the role of DNA sequence in positioning nucleosomes is well understood, a unified framework for studying the impact of transcription remains lacking. Using numerical simulations, we investigated the dependence of nucleosome density profiles on transcription level across multiple species. We found that the low nucleosome affinity of yeast, but not mouse, promoters contributes to the formation of phased nucleosomes arrays for inactive genes. For the active genes, a tug-of-war between two types of remodeling enzymes is essential for reproducing their density profiles. In particular, while ISW2 related enzymes are known to position the +1 nucleosome and align it toward the transcription start site (TSS), enzymes such as ISW1 that use a pair of nucleosomes as their substrate can shift the nucleosome array away from the TSS. Competition between these enzymes results in two types of nucleosome density profiles with well- and ill-positioned +1 nucleosome. Finally, we showed that Pol II assisted histone exchange, if occurring at a fast speed, can abolish the impact of remodeling enzymes. By elucidating the role of individual factors, our study reconciles the seemingly conflicting results on the overall impact of transcription in positioning nucleosomes across species.

## Introduction

Nucleosomes are the fundamental packaging unit of chromatin, comprising 147 base pairs (bp) of DNA wrapped around histone proteins (1). Their formation helps to fit eukaryotic genomes inside the nucleus but also occludes the DNA binding of protein molecules, including regulatory factors and transcriptional machinery (2, 3). The precise position of nucleosomes along the DNA sequence, therefore, can critically impact the function of the genome by regulating its accessibility (4–8). Recent whole-genome sequencing-based studies have indeed revealed the depletion of nucleosomes at many promoter and enhancer regions to accommodate transcription (9, 10). In addition, nucleosome positioning may affect gene expression indirectly by regulating higher-order chromatin organization (11–14). For example, protein molecules such as Cohesin and CTCF have been shown to facilitate chromatin folding and the formation of so-called topologically associated domains (15, 16). These domains promote enhancer-promoter contacts, and their formation relies on the accessibility of CTCF binding sites (17, 18). Underpinning the molecular determinants of nucleosome positioning is, therefore, of fundamental interest and can provide insight into gene regulatory mechanisms.

Since the DNA molecule undergoes substantial distortion when wrapping around histone proteins, its intrinsic, sequence-specific property can impact the stability, and correspondingly position, of the formed nucleosomes (7, 19). Numerous studies have found that nucleosomes preferentially occupy DNA segments that are more susceptible to bending and twisting. They have led to the discovery of periodic dinucleotides (AT and TA) along the nucleosome length (9, 20, 21) and intrinsically stiff poly(dA:dT) tracts at nucleosome-depleted regions (22). Computational models based on such sequence features have been developed to predict *in vivo* nucleosome occupancy (23–25). Accuracy of such predictions can be hampered, however, by the presence of a variety of processes and activities in the nucleus that may overwrite intrinsic positioning signals from the DNA (26–29).

Transcription is one of such processes that can alter the location of nucleosomes via chromatin remodeling and histone eviction (30). Due to the consumption of ATP, the kinetics of these movements does not necessarily satisfy detailed balance, and the resulting nucleosome configurations may conflict with the thermodynamic distribution determined from the DNA sequence alone. The impact of transcription is evident from Figure 1, where average nucleosome density profiles for genes with varying levels of transcriptional activity are shown to exhibit striking differences. In particular, nucleosomes for more active genes (red) appear less ordered in yeast with less pronounced peaks and valleys when compared with inactive ones (blue). However, the opposite trend is observed for mouse, for which clear patterns emerge from a featureless profile as transcription level elevates. The relatively flat density profile of inactive genes is conserved across multi-cellular organisms (31).

**Fig. 1.**
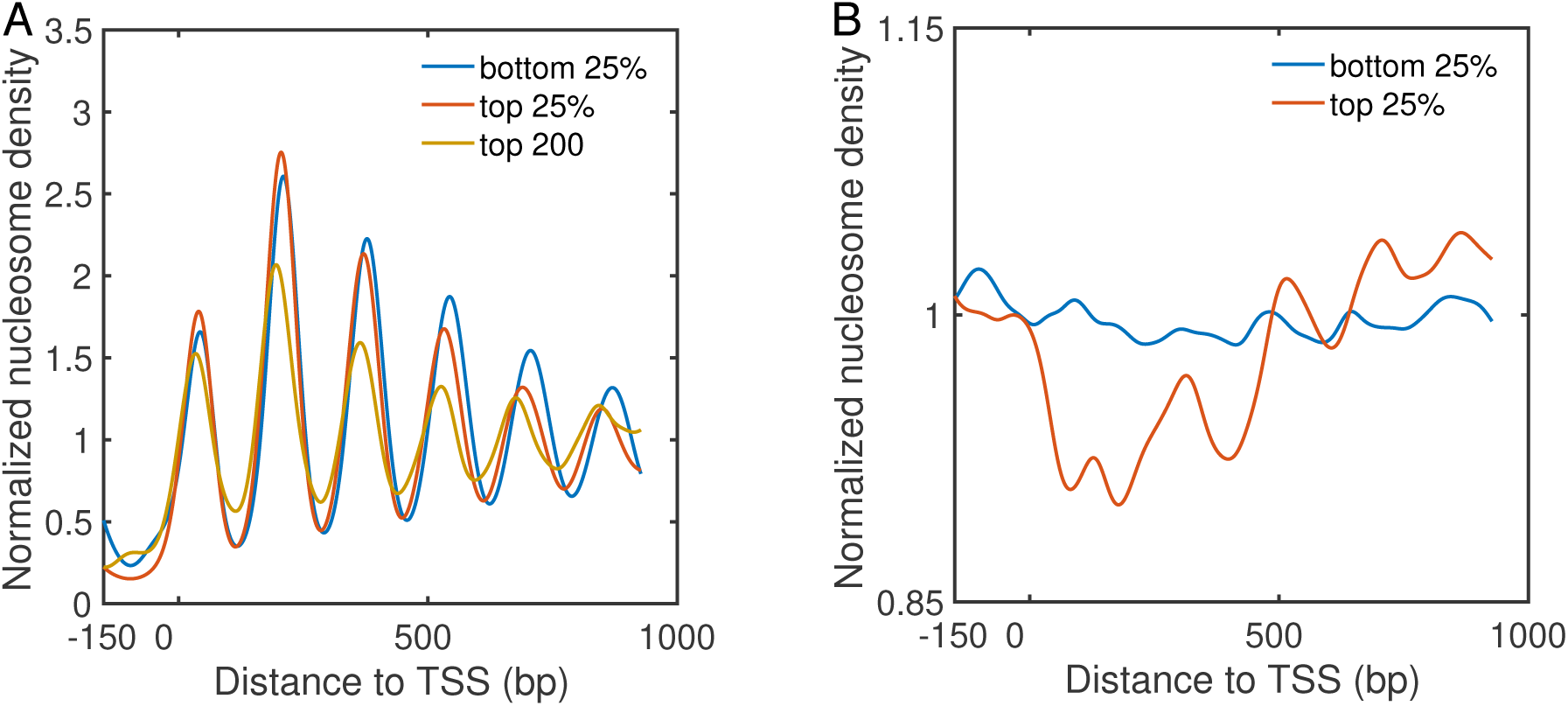
Normalized nucleosome density profiles for *S. cerevisiae* (32) (A) and mouse (31) (B) near TSS. Genes were separated into quartiles depending on levels of transcription activities, with the bottom and top 25% corresponding the most inactive and active genes, respectively. Result for the top 200 most active genes for yeast is also shown to highlight the decrease in amplitude with increased transcriptional activity.

In this paper, we carried out theoretical analysis and numerical simulations to better understand the role of transcription in positioning nucleosomes, and to reconcile the seemingly conflicting trend across species. Our model considers the impact of DNA sequence, transcription factors, chromatin remodeling enzymes, and histone exchange. We found that a tug-of-war between two types of remodeling enzymes explains the observed difference between nucleosome density profiles at varying transcription levels and across species. In particular, remodeling enzymes such as ISW1 that regulate and reduce inter-nucleosome spacing tend to drive the nucleosome array away from the transcription start site (TSS). On the other hand, ISW2-like enzymes help to align nucleosomes towards the TSS. Competition between these enzymes results in two types of density profiles with well- and illpositioned +1 nucleosome. Mixing the two profiles at different levels of population can give rise to results that qualitatively reproduce yeast or mouse data. We further demonstrated that fast kinetics of histone eviction/adsorption, if induced by RNA polymerase (Pol) II elongation, could reduce or abolish the impact of remodeling enzymes. Our study, therefore, provides a unified framework for interpreting the establishment of nucleosome positions inside the nucleus.

## MATERIALS AND METHODS

### Kinetic Model of Nucleosome Positioning

We consider a one-dimensional lattice model to study the positioning of nucleosomes along the DNA sequence (Figure 2). Each lattice site *s* represents a single bp and is assigned with a nucleosome binding energy *V*_*s*_. In most cases, *V*_*s*_ was set to a constant value, such that the nucleosome binding energy of a 147 bp long DNA segment is *V*_*i*_ = –42*k*_*B*_*T* (33), to focus on the impact of remodeling enzymes. When the DNA sequence effect was explicitly considered, we determined *V*_*s*_ using the periodic function of dinucleotides introduced by van Noort and coworkers (24). The length of the lattice is 14700 bp, and the periodic boundary condition was enforced to eliminate any end effect. Nucleosome density was set as 0.88, a typical value found near the gene coding regions in yeast (34).

**Fig. 2.**
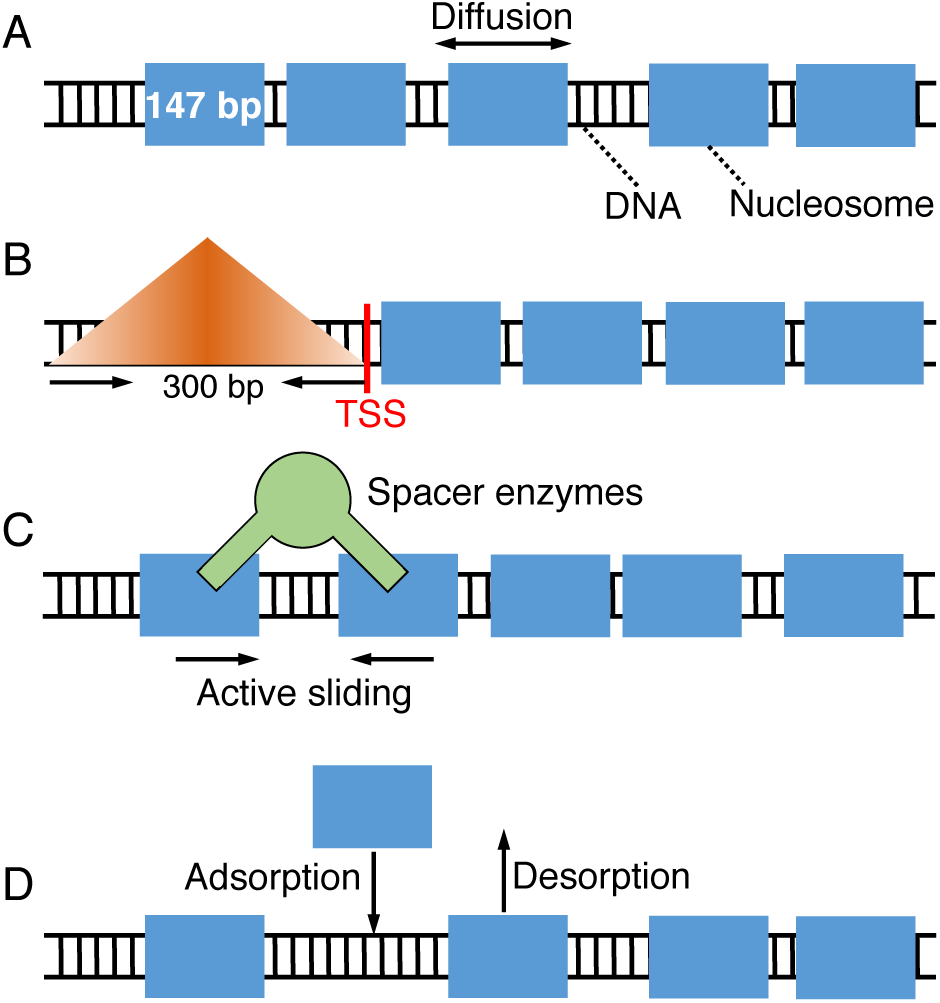
Illustration of the kinetic model used for studying nucleosome positioning that includes thermal diffusion (A), a barrier potential in the promoter region that penalizes nucleosome binding (B), enzyme remodeling (C), and histone exchange (D). The DNA is drawn as a black ladder, and histone proteins are represented as blue rectangles. Remodeling enzymes are colored in green and use a pair of nucleosomes as substrate.

To account for the excluded volume effect, a pair potential was introduced between neighboring nucleosomes *i* and *i* + 1 as

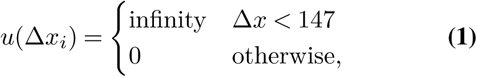

where Δ*x*_*i*_ = *x*_*i*+1_ –*x*_*i*_. *x*_*i*_ corresponds to the dyad position of the *i*-th nucleosome, and by definition, each nucleosome occupies a region of 147 bp in length (1).

Nucleosomes can move along the DNA via diffusive motion with a rate of

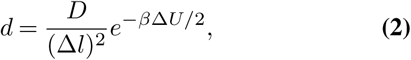

where *D* = 1bp^2^*/*s is the diffusion coefficient and Δ*l* = 1bp is the step size (35). Δ*U* denotes the change of the total energy before and after nucleosome movement. The rate expression was designed such that detailed balance is satisfied (33).

In addition to thermal motions, positions of nucleosomes can be altered by transcription related activities as well (36). In the following, we consider the impact of three major factors related to transcription.

First, transcription factors, preinitiation complex, and Pol II are known to compete with histone proteins to bind gene promoters (37, 38). We incorporated the effect of these proteins as an energetic barrier centered at 150 bp upstream of TSS to penalize nucleosome formation. Similar treatment has been used by Padinhateeri and coworkers to create a nucleosome-free region near TSS (39). As illustrated in Figure 2B, the barrier is symmetric with respect to the center. Its triangular shape allows nucleosomes to occupy the promoter region with a finite probability. The mathematical expression for this barrier potential is provided in the supporting information (SI).

Second, active transcription can recruit remodeling enzymes to alter the position of nucleosomes at the expanse of ATP. While several types of remodeling enzymes have been discovered, here we focus on ISW1-like enzymes that modulate inter-nucleosome spacing and ISW2-like enzymes that adjust the position of the +1 nucleosome (27, 40–45). Following Möbius *et al.* (46), we assumed that spacer enzymes bind to neighboring nucleosomes that are within 332 bp at a rate of 0.16 *s*^−1^ (43, 47), and randomly move one of them toward the other by one bp (Figure 2C). For the positioning enzymes, we modeled their effect with an attractive potential located near TSS.

Finally, transcription of the gene body by Pol II could displace nucleosomes completely off the DNA (28, 48). To account for such disrupt events, we explicitly modeled absorption and desorption of histone proteins with rate expressions

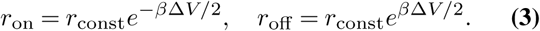

*r*_const_ is the rate constant, and Δ*V* = Δ*U* + *µ* includes both the change in the system’s total energy Δ*U* and a chemical potential *µ*. The impact of Pol II was incorporated into the rate constant and the chemical potential, whose value was tuned to ensure an average system density of 0.88 (see Table S1).

### Details of Stochastic Simulations

We carried out stochastic simulations using the Gillespie algorithm (49) to determine steady-state nucleosome density profiles.

### Simulations without Pol II facilitated histone exchange

In several of the kinetic models explored in the *Results* Section, the effect of histone eviction from Pol II was not explicitly considered. Without remodeling enzymes, these models describe systems with equilibrium statistics since the diffusive dynamics follows detailed balance (Eq. 2). When remodeling enzymes are present, as shown in our previous study (50), the kinetic model can be rigorously mapped onto an effective equilibrium system with renormalized temperature and potential, detailed expressions for which are provided in the SI. For such *one-dimensional* equilibrium or quasi-equilibrium systems, there is a well defined, unique distribution for each model that depends only on inter-nucleosome potentials and DNA sequence. These distributions are independent of the kinetic schemes used in stochastic simulations as long as they satisfy detailed balance. Therefore, for their determination, we simulated only “artificial” absorption and desorption kinetics with rates defined in Eq. 3 and *r*_const_ = 12*s*^−1^. Renormalized potentials were used to determine the change in the system’s total energy Δ*U* if remodeling enzymes were introduced in the kinetic model. The two-dimensional dynamics for histone exchange helps to alleviate the topological constraint and jamming dynamics experienced if the system is restricted to one dimension with all nucleosomes bound to the DNA. It can significantly reduce the computational time needed for convergence. In the large number limit, the statistics of the grand canonical ensemble with histone exchange should be equivalent to that of a system restricted to one dimension with fixed nucleosome number. In Figure S1, we showed that, for the system size considered here, the fluctuation in nucleosome number is small and has minimal impact on the resulting density profile.

We carried out 200 independent simulations for kinetic models lacking DNA sequence specificity. To investigate the impact of DNA sequences, we also separately carried out 1000 simulations for both yeast and mouse. Each one of these simulations incorporates a nucleosome binding affinity profile predicted from the sequence of an inactive gene. All simulations lasted for 5000 seconds and were initialized with over 80 nucleosomes randomly distributed over the lattice. 2500 configurations were recorded every two seconds in each simulation to determine the density profiles.

### Simulations with Pol II facilitated histone eviction

Pol II and remodeling enzymes can evict and assemble nucleosomes during transcription (48, 51). This two-dimensional kinetics for histone exchange defines a steady-state distribution consistent with the reaction rates defined in Eq. 3. The one-dimensional spacer enzymes, by themselves, can give rise to another steady-state distribution that depends on enzyme kinetics. If the rate expressions for histone exchange were modified to account for the effective interaction induced by enzymes, the two steady-state distributions are consistent with each other. This consistency inspired our use of artificial exchange kinetics to accelerate computer simulations mentioned above. Biologically, however, histone exchange rates most likely do not depend on spacer enzymes, and the two distributions will be in conflict.

To rigorously account for the impact of both kinetics, we performed stochastic simulations that explicitly include diffusion, enzyme remodeling, and histone eviction and absorption as well. A total of 2500 independent 5 × 10^5^-second-long simulations were performed. Only 200 configurations in the last 400 seconds of each simulation were collected with an equal time interval to compute the density profiles. These simulations were again initialized with randomly placed nucleosomes over the lattice.

### Data Processing

Genome-wide mappings of nucleosome positions obtained with a chemical mapping method are available for *S. cerevisiae* and mouse in the NCBI database with accession number GSE36063 and GSE82127. Compared to the micrococcal nuclease digestion, followed by high-throughput sequencing (MNase-seq) (29), the chemical mapping approach is affected less by sequence preference or nucleosome unwrapping and can provide base pair resolution of nucleosome center positions (31, 32). We point out, however, that the qualitative trend shown in Figure 1A has been observed for data obtained with MNase-seq as well (30).

To determine the transcription level of individual genes, we downloaded RNA-seq data using accession number GSE52086 for yeast (52) and GSE82127 for mouse (31). DNA sequences surrounding TSS were extracted from the Eukaryotic Promoter Database (53, 54) based on the ID provided in the RNA-seq data.

## RESULTS

### DNA sequence contributes to the barrier for forming phased nucleosome array

A striking difference between yeast and mouse is their distinct nucleosome density profiles for genes with minimal transcription activity (blue lines in Figure 1). While for yeast, these genes exhibit oscillatory patterns with well-positioned nucleosomes, the corresponding curve for mouse is relatively flat with no significant features. Given their low level of transcription, we wondered whether contributions from DNA sequences could explain nucleosome distributions in these genes.

We extracted the sequences surrounding TSS for 1000 genes with the lowest transcription level from yeast and mouse genome. Using a model introduced by van Noort and coworkers that quantifies nucleosome occupancy based on a periodic function of dinucleotides (24), we determined the nucleosome affinity profile for each DNA segment. As shown in Figure 3A, the average affinity for yeast genes quantified in terms of binding energy peaks at promoters located on the left of TSS. Promoters of *S. cerevisiae* are, therefore, inherently nucleosome repelling. Mouse genes, on the other hand, exhibit the opposite trend, with the same region being most favorable for nucleosome formation. The difference in promoters’ nucleosome affinity is particularly interesting in light of the statistical positioning model (10, 55), which argues that the presence of a repulsive potential could create nucleosome-free regions and align downstream nucleosomes. To more directly evaluate the impact of DNA sequences, we carried out simulations for each inactive gene to determine their average density profiles using the predicted sequence-specific nucleosome affinity. Details for these simulations are provided in the *Materials and Methods*. As shown in Figure 3B, it is evident that for yeast but not mouse, there is a depletion of nucleosomes on the left side of TSS. This depletion gives rise to +1 and +2 nucleosomes with well-defined positions, though the peaks do not differ significantly from the results for mouse genes.

**Fig. 3.**
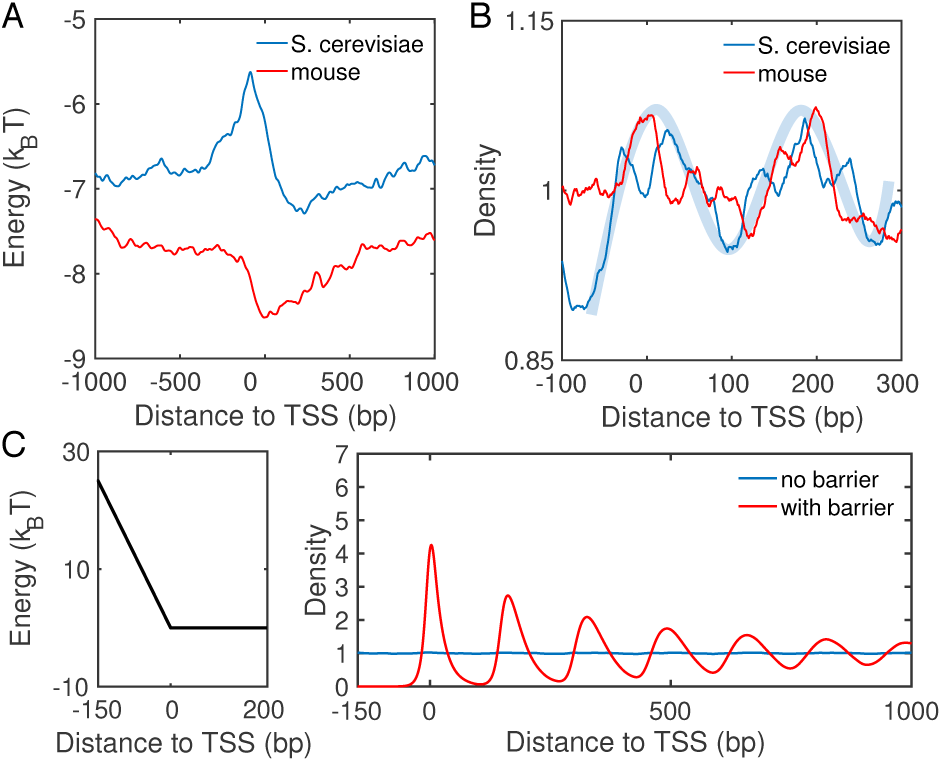
Intrinsic differences in the nucleosome affinity of promoter sequences contribute to the formation of phased nucleosome arrays in yeast, but not mouse, inactive genes. (A) Average nucleosome affinity computed for the 1000 genes with lowest transcription level for yeast (blue) and mouse (red). (B) Nucleosome density profiles for the corresponding sequence-specific affinity shown in part A. The shaded blue curve is shown as a guide for the eye. (C) Nucleosome density profiles for two kinetics models that incorporate a barrier potential in the promoter region or not. An illustration of the barrier is shown on the left.

The less prominent features seen in simulated density profiles can be attributed to the relatively small fluctuation in predicted nucleosome affinity. Additional transcription factors and remodeling enzymes, however, could take advantage of the weakened affinity to occupy promoters, further driving the depletion of nucleosomes and effectively raising the barrier height (37). Without over complicating the model, we incorporated the effect of these proteins with a repulsive potential. As shown in Figure 3C, it similarly increases from TSS to the center of the promoter region (−150 bp) as in yeast affinity profile but with a larger slope. Simulations carried out with this promoter potential resulted in a density profile with clear oscillatory patterns and amplitudes comparable to those seen in experiments. Therefore, the intrinsic property of yeast promoter sequences and the binding of additional protein molecules, or a lack thereof, help to create the nucleosome density profiles of inactive genes.

### Spacer enzymes induce nucleosome condensation and ill-positioned +1 nucleosome

Having resolved the difference between yeast and mouse inactive genes, we next focus on the impact of transcription on nucleosome occupancy in mouse. Figure 1B suggests that as the transcription level increases, nucleosomes become more aligned, as evidenced by the emergence of peaks and valleys. This change could arise from the establishment of nucleosome-free regions at gene promoters to accommodate the arrival of the transcription machinery. Two additional features of the density profile cannot be readily explained by the statistical positioning model, however. First, compared to the curves for yeast (Figure 1A) and from simulations (Figure 3C), the +1 nucleosome in mouse shows a much lower occupancy. Second, the spacing between nucleosomes decreases with the increase of transcriptional activity (Figure S2B). Here nucleosome spacing is measured as the distance between two neighboring peaks. Its decrease has also been confirmed in a recent single-cell study that directly measured the distance between nucleosomes from the same DNA molecule (56). In the following, we explore mechanisms in addition to statistical positioning that can explain these two features.

We note that the decrease of inter-nucleosome spacing upon transcription is indeed a conserved phenomenon and can be readily seen from the yeast profiles as well. In addition, nanopore sequencing of long DNA segments that contain multiple nucleosomes has confirmed the same trend in Drosophila (57). A possible explanation for the spacing change is the recruitment of remodeling enzymes, including ISW1 and Chd1, to actively transcribed genes. These enzymes use a pair of nucleosomes as their substrate and act as rulers to adjust the length of the linker DNA (27, 40, 41, 58). Numerical simulations have confirmed their impact on inter-nucleosome distances via examining the so-called radial distribution profile (46, 50, 59). The impact of these spacer enzymes on nucleosome density profiles near TSS remains unclear, however.

We carried out simulations to study the distribution of nucleosomes with the presence of spacer enzymes and a barrier potential in the promoter region to approximate the impact of DNA sequence. As detailed in the *Materials and Methods* section, these enzymes bind with a pair of neighboring nucleosomes and move them closer by one bp at every step. The rate of such remodeling steps is independent of the underlying energy landscape, and the enzymes break the detailed balance. Using a theory developed by us (50), we mapped the non-equilibrium model with enzymes onto an equivalent and renormalized equilibrium system with effective, attractive interactions between nucleosomes. To determine the distribution of nucleosomes for this effective equilibrium system, we used artificial dynamics that significantly reduces the computational time needed for statistical convergence. As shown in Figure S3, the average distance between neighboring nucleosomes indeed decreases upon the introduction of enzymes. To our surprise, however, the density profile resembles that for active genes from mouse, with depletion of nucleosomes near TSS (see Figure 4A). It is in striking difference from the density profile for a model with only the barrier potential (Figure 3C). The well-positioned +1 nucleosome disappears, and nucleosome density shifts towards downstream regions. Examining the simulated nucleosome arrays revealed a wide range of configurations with both well- and ill-positioned +1 nucleosome. We first ordered the nucleosome arrays along the *y*-axis based on the position of the first nucleosome and computed the corresponding local nucleosome density profiles. The results are shown in the left panel of Figure 4B, with representative, traditional one-dimensional profiles presented on the right. The top configurations exhibit a well-defined +1 nucleosome, and the corresponding density profile resembles that of a statistical positioning model shown in Figure 3C. For many of the configurations near the bottom of the plot, the +1 nucleosome shifts away from the TSS, giving rise to a wide nucleosome free region. A mixture of these configurations with varying +1 nucleosome positions results in the final profile shown in Figure 4A.

**Fig. 4.**
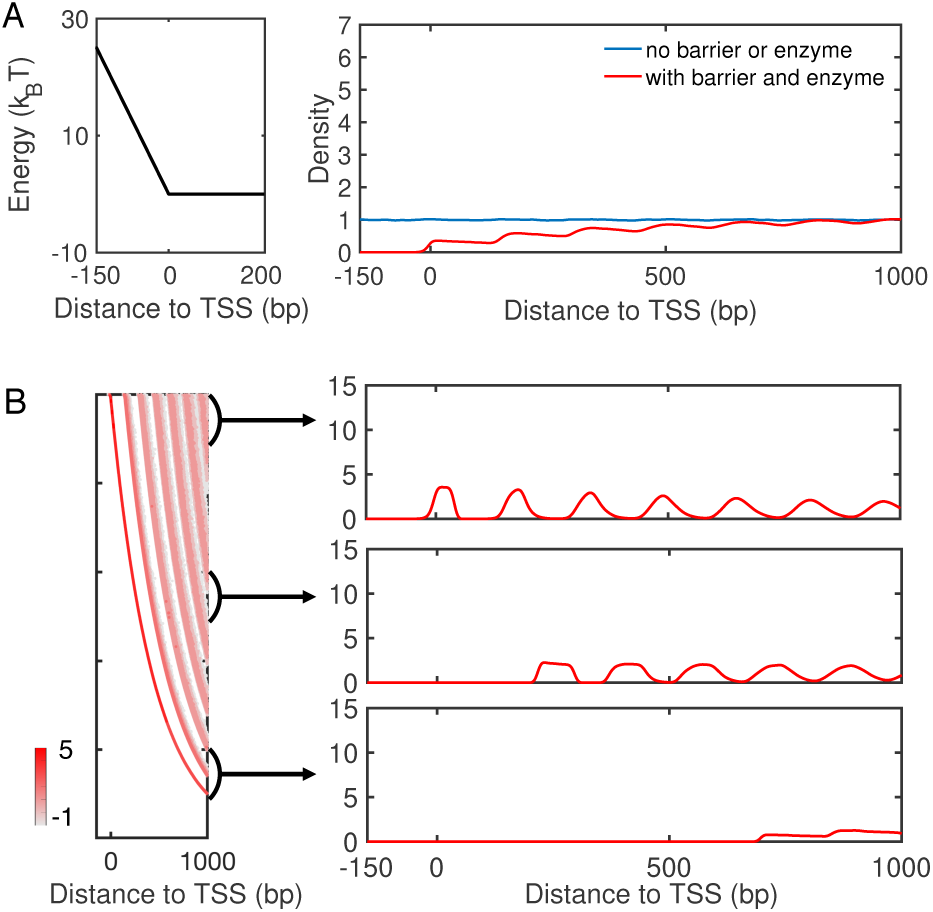
Spacer enzymes drive nucleosome condensation and the formation of ill-positioned +1 nucleosome. (A) Simulated nucleosome density profile using a model that includes a barrier potential (left) and spacer enzymes reproduce results for mouse active genes. (B) Nucleosome configurations exhibit a wide distribution of positions for the +1 nucleosome. A scatter plot for the density distribution of all simulated configurations ordered by the position of the first nucleosome is shown on the left. The right panel presents three example one-dimensional profiles in which the +1 nucleosome gradually shifts away from the TSS.

The inclusion of spacer enzymes can, therefore, impact both inter-nucleosome distances and the position of the +1 nucleosome. Without these enzymes, nucleosomes will occupy all accessible DNA regions while staying as far apart from each other as possible to maximize entropy. This tendency for an equal partition of the DNA is the essence of the statistical positioning model. It will ensure the confinement of the +1 nucleosome in a narrow region between the TSS and the +2 nucleosome. On the other hand, spacer enzymes introduce effective attraction between nucleosomes and cause them to aggregate rather than staying farther apart (50). The entire array of nucleosomes now behaves as a single entity, and individual nucleosomes are no longer uniformly distributed across the genome. The free, collective movement of the entire nucleosome array with respect to the TSS, again driven by entropy, will result in ill-positioned +1 nucleosome.

### A mixture of profiles with well- and ill-positioned +1 nucleosome reproduces yeast results

The presence of a barrier potential and spacer enzymes leads to the formation of two types of nucleosome density profiles with well- and ill-positioned +1 nucleosome. A mixture of the two types qualitatively reproduces the experimental results for active mouse genes. We next investigated whether the same mixture but with different levels of population can explain yeast nucleosome density profiles.

The more pronounced patterns seen in yeast profiles suggest that configurations with well-positioned +1 nucleosome should dominate. We note that many proteins, including remodeling enzyme ISW2, are known to align nucleosomes toward the TSS. To mimic the impact of these molecules, we introduced an additional attractive potential between 150 and 180 bp from TSS. We chose this region rather than the one immediately following the promoter to reproduce the higher density for the +2 nucleosome. This site can indeed be more favorable for nucleosome formation as the binding of transcription factors is known to extend beyond the promoter to compete with nucleosome formation at downstream sites (37).

As shown in Figure 5, the new potential succeeds in attracting nucleosomes to the TSS, and the resulting density profile (red) now resembles those from yeast. We note that as transcription activity decreases, enzymes will be recruited less to the genes, and their effective remodeling rate will be smaller. Slower spacer enzymes with a rate of *k* = 0.08*s*^−1^ compete less effectively with the positioning enzymes, and the relative population of configurations with ill-positioned +1 nucleosome decreases. Correspondingly, the nucleosome profile exhibits higher peaks (blue) than that for more active genes with a faster enzyme rate, consistent with the dependence on transcription activity seen in experimental results (Figure 1A).

**Fig. 5.**
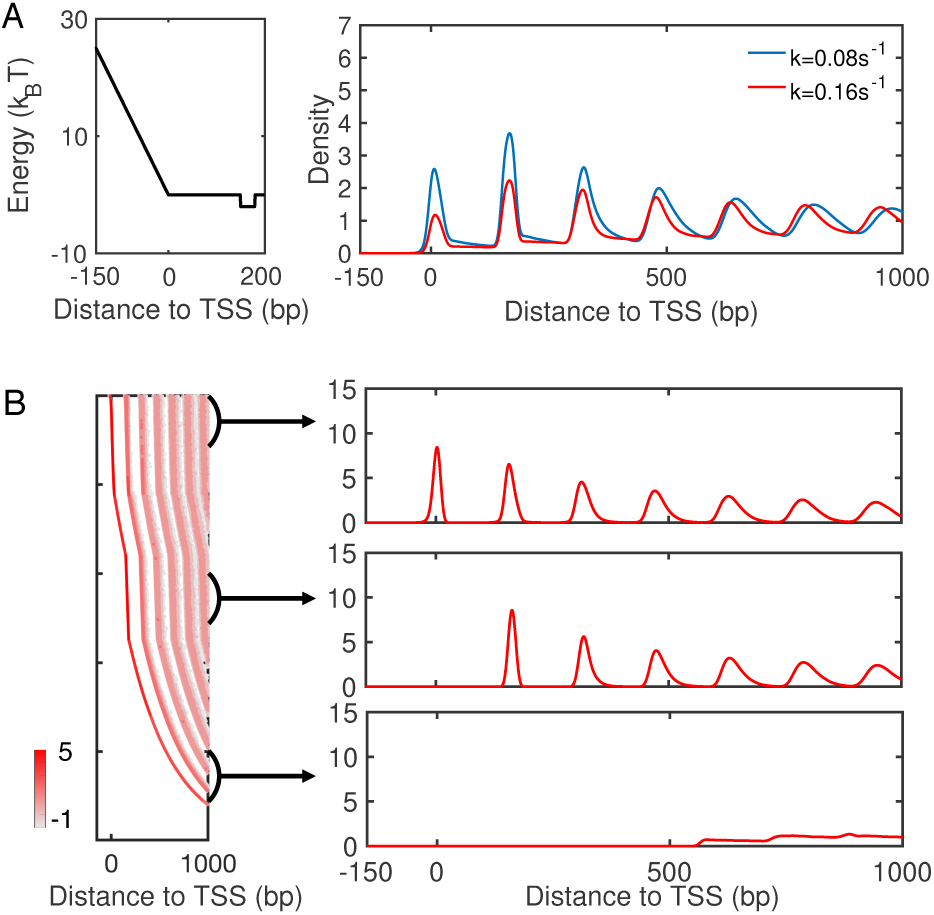
A mixture of configurations with well- and ill-positioned +1 nucleosome reproduce yeast density profiles. (A) Nucleosome density profiles determined with the presence of an attractive potential of −2*k*_*B*_*T* introduced to the region following the promoter (left). Comparison between the two density profiles with varying rate for spacer enzymes confirms that more active genes with higher enzyme rates exhibit lower peaks. (B) Scatter plot for the density distribution of all simulated configurations ordered by the position of the +1 nucleosome (left), with example one-dimensional profiles shown on the right.

### Fast histone exchange abolishes the impact of spacer enzymes

For highly transcribed genes, in addition to remodeling enzymes, Pol II could impact the positioning of nucleosomes as well. As it elongates along the DNA, Pol II could cause partial or complete loss of histone proteins (28, 48, 51, 60). In the following, we investigate the impact of Pol II induced histone eviction on nucleosome density profiles.

Specifically, we carried out stochastic simulations that explicitly model nucleosome diffusion, enzyme remodeling, and histone eviction and absorption. We assumed that the eviction and absorption rates depend only on inter-nucleosome and nucleosome-DNA interactions and are independent of remodeling enzymes (see *Materials and Methods*). The ratio of the two was tuned to ensure a density of approximately 0.88.

The basal rate constant *r*_const_ = 0.1*s*^−1^ was estimated from the transcription rate 1 min−1 for the most active genes (61) with the assumption of full eviction for all nucleosomes.

The resulting nucleosome density profile is shown in Figure 6 (purple). It differs significantly from the one obtained from a kinetic model that only included remodeling enzymes and a barrier potential (red), which was also shown in Figure 4A. The impact of remodeling enzymes in these simulations is significantly reduced. In particular, a pronounced peak emerges near TSS, and the density profile now traces well the result from a model with only the barrier potential (blue). The decrease of inter-nucleosome spacing cannot be observed in the radial distribution profile either (Figure S4). We found that the impact of histone eviction on the density profile depends on its rate and gradually diminishes as the rate slows down (Figure S5). It is worth noting that a significantly smaller rate (10^−8^*s*^−1^) is needed to reveal the impact of remodeling enzymes on nucleosome spacing. Since a decrease of inter-nucleosome spacing is readily seen in experimental nucleosome density profiles, we anticipate that complete eviction of histone octamers to be rare (28, 34, 62, 63). The competition between Pol II and spacer enzymes on positioning nucleosomes can be understood as following. The diffusive dynamics driven by thermal motions defines an equilibrium distribution of nucleosomes along the DNA. This distribution depends both on inter-nucleosome and nucleosome-DNA interactions. Spacer enzymes modify this distribution by introducing an effective attractive potential between nucleosomes. The two-dimensional dynamics of histone exchange can give rise to, yet, another steady-state distribution. Unless slowed down substantially, histone exchange can lead to faster relaxation kinetics when compared with nucleosome movements restricted to one dimension. It will essentially overwrite any impact caused by spacer enzymes or diffusion on nucleosome distribution. If the rates for histone eviction and adsorption satisfy detailed balance with regard to the normal potential for nucleosome-DNA and inter-nucleosome interactions, the steady-state distribution determined from histone exchange kinetics should be consistent to the equilibrium distribution obtained from pure diffusion. This consistency explains the agreement between purple and blue lines seen in Figure 6A. On the other hand, if the two rates were modified to account for the effective interaction potential induced by spacer enzymes, the steadystate distribution will reproduce the one dictated by spacer enzymes. Such kinetics, though less meaningful biologically, can prove beneficial for reducing the computational cost of stochastic simulations (see *Materials and Methods* for more discussions).

**Fig. 6.**
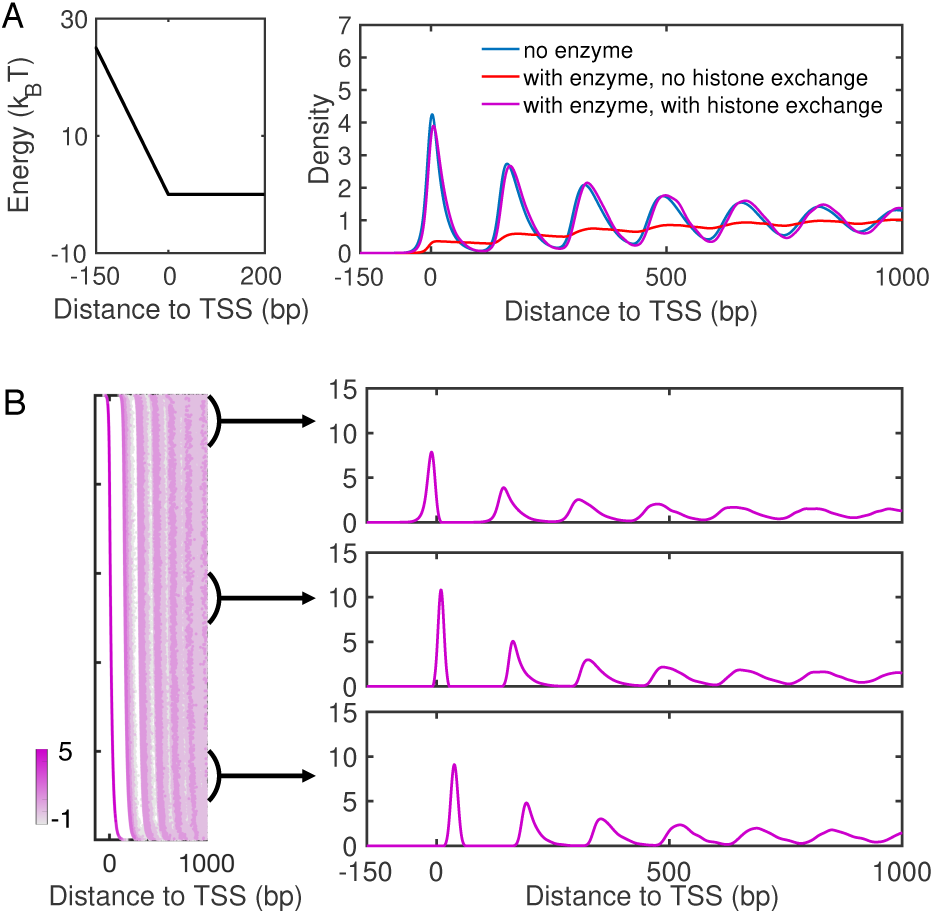
Impact of histone exchange kinetics on nucleosome density profiles. (A) Nucleosome density profiles determined from kinetic models that only includes a barrier potential (blue), that includes both a barrier potential and spacer enzymes (red), and that includes a barrier potential, spacer enzymes and histone exchange (purple). The red curve is identical to the one shown in Figure 4A. An illustration of the barrier potential is shown on the left. (B) Scatter plot for the density distribution of all simulated configurations ordered by the position of the +1 nucleosome (left), with example one-dimensional profiles shown on the right.

## CONCLUSIONS AND DISCUSSION

In this paper, we investigated the impact of transcription on nucleosome positioning. By partitioning genes based on their transcriptional activity, we determined the corresponding nucleosome density profiles for both yeast and mouse. A striking difference for inactive genes was observed between the two species. Similar featureless profiles as that from mouse have been observed for inactive genes from Drosophila(57) and human(64) as well. Analyzing the nucleosome binding affinity of DNA sequences suggests that while yeast promoters are nucleosome repelling, the opposite holds true for mouse promoters. This difference could contribute to the formation of phased nucleosome arrays in yeast, but not mouse, via the statistical positioning mechanism. The nucleosome attracting promoters appear to be a rule of multi-cellular organisms rather than an exception in mouse, as shown in prior studies (23, 65, 66). They might function to suppress the expression of certain genes crucial for cell differentiation (66).

We further carried out stochastic simulations to study the variation of nucleosome density profiles as the transcriptional activity elevates. We discovered that a tug-of-war between two types of enzymes is the key to rationalize the observed trends. In particular, enzymes that use a pair of nucleosomes as substrate, including Chd1 and ISW1, can induce nucleosome condensation and tend to shift the nucleosome array away from TSS, giving rise to density profiles with ill-positioned +1 nucleosome. Enzymes such as ISW2, on the other hand, can counteract this effect and align the +1 nucleosome back to TSS. A combination of density profiles with well- and ill-positioned +1 nucleosome can qualitatively reproduce *in vivo* results from both yeast and mouse. A significant difference between the two profiles is the length of the nucleosome-free region, and genome-wide nucleosome positioning profiles indeed support the presence of both narrow and wide promoter regions in mouse (56, 67) and yeast (68).

## ACKNOWLEDGEMENTS

This work was supported by the National Institutes of Health (Grant 1R35GM133580-01). The authors thank John van Noort for sharing the software for nucleosome affinity calculation.

